# Patterns of regulatory divergence and gene expression in hybrids are associated with molecular evolution in species undergoing gene flow

**DOI:** 10.1101/2021.11.14.468549

**Authors:** Fernando Díaz, Jason Wolf, Reinaldo de Brito

## Abstract

The extent to which hybridization disrupts a gene’s pattern of expression likely governs its propensity for introgression, while its extent of molecular divergence can itself underlie such disruption. Together, these phenomena shape the landscape of sequence and transcriptional divergence across the genome as species diverge. To understand this process, we examine gene expression inheritance, regulatory and molecular divergences in the reproductive transcriptomes of species linked by gene flow. The fruit flies *Anastrepha fraterculus* and *A. obliqua* show evidence of gene flow despite clear evolutionary divergence and incomplete reproductive isolation. We find that their transcriptional patterns are a mosaic between those typically observed within and between allopatric species. Genes showing *transgressive* expression in hybrids or *cis*-regulatory divergence between species are associated with greater molecular divergence. This may reflect pleiotropic constraints that make them more resistant to gene flow or they may be more likely to experience divergent selection. However, while these highly divergent genes are likely to be important contributors to species differences, they are relatively rare. Instead, most differentially regulated genes, including those linked to reproduction, show high degrees of *dominance* in hybrids and *trans*-regulated divergence between species, suggesting widespread genetic compatibility that allowed for the identified introgression. These findings provide insights into how postzygotic isolating mechanisms might evolve in the presence of gene flow: regions showing *cis*-regulatory divergence or *transgressive* expression contribute to reproductive isolation, while regions with *dominant* expression and *trans*-regulatory divergence act as a buffer of hybrid breakdown, facilitating introgression, and leading to a genomic mosaic of expression and sequence divergence.

## Introduction

Although species are often considered to be well-defined evolutionarily independent lineages, there is increasing evidence that there is a continuum of reproductive isolation and, hence, genetic exchange (Mallet et al., 2007; Martin et al., 2013; Harrison and Larson, 2014). As a result, the genomes of otherwise well-defined divergent species are often mosaics composed of regions homogenized by introgressive hybridization (Wu, 2001) and divergent regions that are responsible for the genetic isolation (Harrison and Larson, 2014). Although these “genomic islands” are well documented across different species and their molecular evolution has received considerable attention (Harr, 2006; Mallet et al., 2007; Malinsky et al., 2015; Choi et al., 2020), their causes are still in debate (Kulathinal et al., 2009; Cruickshank and Hahn, 2014; Han et al., 2017). The propensity for introgression and the extent of molecular divergence that shape the pattern of these islands are presumably governed by the array of phenomena that ultimately result in genetic incompatibilities, including transgressive phenotypes, genomic rearrangements, altered protein interactions, patterns of gene expression, and disrupted epigenetic mechanisms (Ortiz-Barrientos et al., 2007; Barbash, 2010; Maheshwari and Barbash, 2011). Individual genes and potentially larger genomic blocks can vary in their contribution to these phenomena, and hence will vary in their propensity for introgression and degree of divergence. Those regions that remain compatible are able to introgress, potentially counteracting divergence by selection and drift (Mallet et al., 2007; Martin and Jiggins, 2017; Nadeau and Kawakami, 2018), whereas selection on regions associated with disrupted expression or phenotypes in hybrids will contribute to maintain or foster divergence.

Many of the phenomena governing species divergence and propensity for introgression are tied to the regulation of gene expression and associated disruption of expression in hybrids, suggesting that the mosaic of divergence can be, at least in part, understood through characterization of patterns of hybrid gene expression (Wittkopp et al., 2004; Signor and Nuzhdin, 2018). Genes in regions showing recent or ongoing gene flow across species would presumably show patterns of expression consistent with genetic compatibility while genes in regions resistant to gene flow would presumably show patterns of expression disruption that could underlie genetic incompatibilities. For example, individual genes (and their regulatory regions) can show sequence divergence that causes *cis*-regulatory differences (Wei and Zhang, 2018), which can lead to allelic imbalance in hybrids (Coolon et al., 2014; Landry et al., 2005; Signor and Nuzhdin, 2018). As a result, *cis-*regulated genes may be resistant to introgression because they maintain their divergent expression profile, while those that are *trans*-regulated may adopt the profile of the "host" genome, which could facilitate introgression. This idea is supported by studies that, at a gross scale, suggest that *cis*-regulatory mechanisms are particularly associated with highly divergent genomes (Wittkopp et al., 2004, 2008; Ortiz-Barrientos et al., 2007; Graze et al., 2012; Bell et al., 2013; Coolon et al., 2014; Meiklejohn et al., 2014; Chen et al., 2015; Signor and Nuzhdin, 2018), while *trans*-regulatory mechanisms are more likely to account for expression divergence between genomes with low divergence, such as intraspecific crosses (Bell et al., 2013; Suvorov et al., 2013; Signor and Nuzhdin, 2018). Most intra and interspecific patterns of gene regulation reflect the broader patterns of regulatory element evolution between species and are typically addressed by studying allopatric species. While the regulation and inheritance of gene expression in hybrids is likely a key factor in shaping the pattern of divergence across genomes, the relationship between these phenomena and molecular divergence in species showing isolation with gene flow remains poorly understood (Graze et al., 2012; Meiklejohn et al., 2014). Available evidence, particularly from closely related *Drosophila* species (Coolon et al., 2014; Meiklejohn et al., 2014; Banho et al., 2021) that regulatory divergence in recently established species represents an intermediate mosaic with respect to that seen within species and between highly divergent species. However, the role of gene flow in shaping the landscape of introgression between recently diverged species is still unresolved (Martin and Jiggins, 2017).

To understand the relationship between regulation and inheritance of gene expression in hybrids and patterns of molecular evolution between species, we investigate these using a pair of neotropical fruit fly species, *Anastrepha fraterculus* and *A. obliqua* that diverged ^~^2.6 Mya and show signatures of divergence with gene flow (Díaz et al., 2018). These species belong to the *fraterculus* group, which harbor some of the most important agricultural pests in South America (Aluja, 1994). Despite the presence of gene flow, these species can easily be distinguished by several phenotypic differences and show differences in ecologically relevant traits such as host preferences and reproductive behavior (Aluja, 1994); (Sivinski et al., 1999); (Aluja et al., 1999). Detected gene flow suggests that prezygotic barriers to reproduction are leaky (Scally et al., 2016; Díaz et al., 2018). Indeed, hybrids between these species can be obtained in the laboratory, but postzygotic incompatibilities follow Haldane’s rule (Selivon et al., 1999; Santos et al., 2001; Rull et al., 2017). Because genes associated with reproductive isolation are likely to evolve rapidly despite gene flow, making their patterns of expression divergence more detectable from reproductive tissues (Andrés et al., 2008), we focus on the reproductive transcriptomes of the parental species and their F_1_ hybrids. Using these we characterize patterns of expression inheritance (e.g., additive/transgressive) and regulatory divergence (e.g., *cis*/*trans*) and relate these to genome-wide patterns of molecular evolution and gene flow to provide insights into how postzygotic isolating mechanisms might evolve in the presence of gene flow while buffering the reproductive breakdown that facilitates introgression.

## Methods

### Study population

Crosses were derived from established lab populations of *A. fraterculus* and *A. obliqua* originally collected from fruits of hostplants in Midwest (16° 41’ 58’’ S, 49° 16’ 35’’ W) and Southeastern (22° 01’ 03’’ S, 47° 53’ 27’’ W) regions of Brazil respectively. Field collected flies were identified using wing, ovipositor, and other morphological markers following identification keys available in (Norrbom et al., 2012). These lab populations were maintained in the Population Genetics and Evolution Lab at the Federal University of São Carlos (Brazil) for over two years in a controlled environment room at 26 °C (60–90% humidity) and natural photoperiod before the experiment. Mango (*Mangifera indica* L.) fruits were used for oviposition and larval development while adults were fed on a mixture of hydrolyzed protein, vitamins, and sucrose. Populations were maintained by sampling over 100 mating pairs of adults to generate non-overlapping generations and reduce inbreeding.

Samples from the parental species were derived from individual intraspecific crosses (individually paired *♀ A. fraterculus* x *♂ A. fraterculus* and *♀ A. obliqua* x *♂ A. obliqua*) while reciprocal F1 hybrids were derived from interspecific crosses (individually paired *♀ A. fraterculus* x *♂ A. obliqua* and *♀ A. obliqua* x *♂ A. fraterculus*). Because the *♀ A. obliqua* x *♂ A. fraterculus* cross only produces female progeny (Santos et al., 2001), our analysis is restricted to three of the four classes of hybrids. To indicate the direction of the cross we refer to the reciprocal hybrids as OF and FO (F = *A. fraterculus* and O = *A. obliqua*, where the female parent is listed first). Entire male and female reproductive tissues were obtained from 10 days-old mature virgin progeny from each of the crosses. These tissues were chosen because they are likely to express genes that play important roles in species differences as they tend to evolve more rapidly than background genome (Andrés et al., 2013). Groups of five specimens were pooled for each sample and three replicates were generated per cross, which generated six samples for each parental species and FO hybrids and three for OF hybrids (only females), for a total of 21 samples. All samples were kept on Trizol reagent at −80 °C until RNA extractions.

### RNA extraction, cDNA library construction, and sequencing

Total RNA was extracted from pooled samples using the Trizol-chloroform protocol (Chomczynski and Mackey, 1995). RNA quality was visually inspected by agarose gel electrophoresis and quantified using both a Qubit fluorometer and Nanodrop spectrophotometer. cDNA libraries were created using Illumina TruSeq Stranded mRNA Sample Prep LS Protocol according to the manufacturer’s instructions. Libraries were sequenced at the Laboratory of Functional Genomics Applied to Agriculture and Agri-energy, ESALQ-USP, Brazil using the HiSeq SBS v4 High Output Kit on Illumina platform flow cells with runs of 2 × 100 bp paired-end reads. Illumina’s HiSeq Control Software and CASAVA v1.8.2 software (Illumina, Inc.), were used for base calling and sample demultiplexing.

### Sequence trimming and assembly

Reads were trimmed for quality and adapter sequences were removed using a minimum quality base of Q=20 and minimum read length of 50bp using Trimmomatic (Bolger et al., 2014). Because there is no reference genome for either species, *de novo* assembly was performed using the software Trinity (Grabherr et al., 2013) with default parameters.

### Improving transcriptional assemblies before mapping

*De novo* assemblers as implemented in Trinity often generate hundreds of thousands of sequences resulting in a large number of isoforms grouped in components, often with numerous redundancies and chimeric sequences (Yang and Smith, 2013). Trinity may generate several sequences that correspond to the same gene (due to alternative splicing) or independent sequences from the same gene as well as combined genes or chimeras. Such complications are maximized when there is increased heterogeneity in the samples, such as when combining samples from different individuals and species (Yang and Smith, 2013). To minimize these issues, we performed separated assemblies for male and female reproductive tissues. We then used an assembly cleaning strategy based on the software set Trinity-Bowtie2-RSEM-CD-HIT-EST, to reduce redundant and chimeric sequences, while maximizing the number of truly representative isoforms. This strategy has been empirically demonstrated to show superior results when compared with alternative approaches, improving the accuracy of downstream analyses (Yang and Smith, 2013). We used Trinity utilities to filter the assembled transcriptomes by abundance, and mapping reads back to their assemblies using Bowtie2 (Langmead et al., 2009). A read count matrix was generated using RSEM and only isoforms with the highest percentage of abundance within each component were retained as representative based on read counts after TMM normalization. Then, we reduced the remaining redundancy using CD-HIT-EST with a sequence identity cutoff set to 0.98 to collapse highly similar components within assemblies.

### Allele-specific expression

Analyses of allele-specific expression (ASE) rely on the identification of the parental origin of reads sequenced in hybrid samples. For this, we identified fixed SNP differences between the two populations. Reads coming from the allele more similar to the reference can potentially map with higher probability or quality than reads coming from the non-reference allele, leading to inaccurate estimates of ASE. To avoid this mapping bias, we accounted for polymorphic variation in the reference using GSNAP (Wu and Nacu, 2010), which allows SNP-tolerant alignment in mapping. This method allows SNP information to be introduced into the reference, which allows reads from two different species to be mapped to a “common diploid *Anastrepha* reference” that includes all the divergent SNPs. This SNP-tolerant strategy has been shown to better account for allele mapping bias than alternatives strategies such as SNP masking or reciprocal best hits (Satya et al., 2012).

SNP calling and SNP-tolerant mapping were carried out using modules implemented in the Allim pipeline (Pandey et al., 2013). For this, only one of the initial assemblies (*e.g., A. obliqua*) was retained for each profile, and this assembly was used as a template to generate the “common diploid *Anastrepha* reference” used in further analyses. Allim can automate multiple rounds of SNP identification and GSNAP mapping (with or without the SNP-tolerant option). After mapping the reads from both species to the selected reference assembly, divergent SNPs were identified. Polymorphism information was then used in a subsequent round of SNP-tolerant mapping. SNPs were identified using SAMtools (Li et al., 2009) with the mpileup option, which was then used to call the SNPs through Bayesian inference in bcftools (Li et al., 2009). Because the SNP calling was performed to identify fixed SNPs between species, only SNPs where the species were fixed for different alleles were considered. This allows reads to be unambiguously assigned to one of the parental genotypes. This information was then incorporated into the mapping for an SNP-tolerant alignment via GSNAP. Allim was also used to assess the quality of the common reference by testing for any remaining bias using simulated reads. For this, both references generated were used to simulate the same number of reads for each polymorphic site, and then simulated reads were mapped back to the “common diploid *Anastrepha* genome”.

### Expression divergence and allelic imbalance

We created a read count matrix for the parental species and hybrids using the reads that were unambiguously assigned to their specific parental origin. Count data were rounded to the nearest integer to satisfy the requirements of downstream statistical tests (e.g., negative binomial). The read count matrix was filtered for a minimum count cutoff of 3 cpm for each parental species and hybrids over at least two out of three replicates per comparable group. All zero values were then adjusted to one to satisfy the binomial tests used below for *cis* and *trans* classifications since positive integers are required. All expression analyses were performed using the R package *edgeR* (Robinson et al., 2009) after TMM library normalization. Normalized counts were analyzed by Generalized Linear Models accounting for the negative binomial variable of read counts in the case of gene expression as well as binomial variable for ASE in hybrids, followed by analyses of expression divergence (ED) between species, modes of inheritance (e.g., additive / non-additive) and regulatory divergence (e.g., *cis*/*trans*). An FDR correction (Benjamini and Hochberg, 1995) using a global α = 0.05 for multiple comparisons as well as a log_2_-fold-change threshold of > 1.25 was applied to all *P*-values.

### Inheritance modes of gene expression

Modes of inheritance were investigated by comparing global expression levels of a given gene between the parental species *Log*_2_*(P*_1_*/P*_2_) and between hybrids and each parental species: *Log*_2_*(P*_1_*/F*_1_) and *Log*_2_*(P*_2_*/F*_1_) while ignoring allelic information (total gene expression levels in hybrids reflect the sum of reads mapped to both parental alleles). A negative-binomial GLM analysis was used to evaluate pairwise comparisons of gene expression between hybrids and their parental species. Based on these comparisons, each transcript was classified according to a commonly used system to describe *additive* and *non-additive* modes of gene expression (McManus et al., 2010); (Bell et al., 2013). Non-differential transcripts between parents and hybrids were classified as conserved. Transcripts for which expression in the hybrid is not significantly different from one of the parents were considered as *dominant* for that parent. Transcripts for which hybrid expression was not similar to either parent but is within the parental range were classified as *additive*, while transcripts for which expression in the hybrid is either above or below parental range were considered as transgressive (*overdominant* and *underdominant* respectively).

### Cis/Trans regulatory divergence

Gene expression can be regulated by *cis*-acting or *trans*-acting effects (Figure 1). The contribution of regulatory effects on gene expression was investigated by comparing the extent of allele imbalance in hybrids (i.e., the relative expression of allele *A*1 derived from species 1 and allele *A*_2_ from species 2):

**Figure 1.**
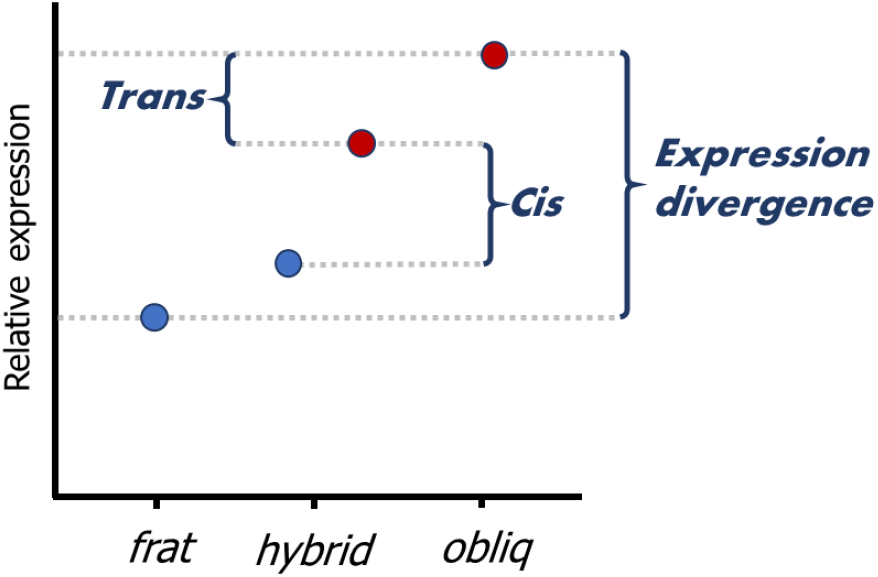
Schematic representation of mechanisms of regulatory divergence (*cis* and *trans*) between *A. fraterculus* (*frat*) and *A. obliqua* (*obliq*). Because alleles in hybrids are exposed to the same genetic background, allele-specific expression (ASE) in hybrids represent the level of expression divergence due to *cis*-acting regulation, while the *trans-*acting component can be extracted from total expression divergence between species in comparison with the ASE in hybrids. The *cis* index is then calculated from the relative contribution of *cis* vs *trans* effects to total expression divergence [Figure taken and modified form (Signor and Nuzhdin, 2018)].

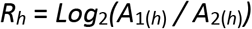

and the relative expression of these alleles when homozygous in the parental species (Haerty and Singh, 2006); (McManus et al., 2010); (Bell et al., 2013):

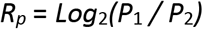

In the genetic regulatory background of the hybrid, both parental alleles are exposed to the same *trans*-effects, which means that any allele imbalance will be the result of a *cis*-regulatory effect (hence, *R*_*h*_ represents the *cis*-regulatory effect, where the allele present at a locus is responsible for variation in its own expression). Consequently, any deviation on the degree of expression divergence between parents and the degree of allele imbalance in hybrids indicates the occurrence of *trans*-regulatory effects. Hence, the trans effect is given by:

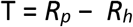

A negative-binomial GLM analysis was used to evaluate significant *cis*-effects, based on allelic imbalance in the hybrids. *Trans*-effects were identified through a binomial GLM comparing the ratios of ASE between hybrids (*R*_*h*_) and parents (*R*_*p*_). Thus, non-differential ratios are evidence of *cis*-only regulatory divergence, while any significant deviation between ratios is evidence of additional *trans*-effects favoring the expression of one allele.

Based on the extent and direction of expression change, we classified transcripts in seven different patterns. Transcripts were classified as *conserved* if there was no significant differential expression between parents, between alleles in hybrids, and between their ratios. Transcripts show *cis*-only regulation when there is significant differential expression between parents, and that pattern is retained for the alleles in hybrids, but no significant differences between their ratios. Transcripts show *trans*-only regulation when there is significant differential expression between parents, but not between alleles in hybrids, with significant differences between their ratios. Transcripts may also show a mixture of these, with significant differential expression between parents, between alleles in hybrids, and significant differences between their ratios. In the case where there is an expression difference between parental and hybrid alleles that goes in direction of the same allele, transcripts are classified as *cis + trans*, whereas when the expression differences are biased towards different alleles, transcripts are classified as *cis * trans*. Transcripts are classified as *compensatory* when there is no significant differential expression between parents, but there are significant differences between alleles in hybrids and significant differences between their ratios. In this case, opposite changes between *cis* and *trans*-effects compensate each other, resulting in no expression differences between species. Finally, transcripts are classified as *ambiguous* when expression patterns did not follow any clear expectations according to these criteria.

### The cis index

To characterize the overall pattern of regulation, we quantified the relative effect of *cis*-acting vs *trans*-acting mechanisms for each transcript. For this, we created an index that measures the size of the *cis* component relative to the total *cis* and *trans* effects:

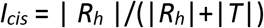

This index was used to investigate the relative role of these mechanisms concerning expression inheritance and molecular and expression divergences.

### Molecular divergence between species

We estimated molecular divergence between species using the ASE dataset. For this, an additional SNP-tolerant mapping via GSNAP using only parental samples was performed. SNPs between species were identified using SAMtools (Li et al., 2009) with the *mpileup* option, and then allele frequencies were obtained using Popoolation II software with a minimum quality base of Q=20 and minimum allele frequency MAF = 2%. We then calculated the interspecific differentiation index (*d*) as the absolute difference in allele frequencies between species (|*frat - obliq*|) for each SNP, and then estimated the molecular divergence as the percentage of fixed variation per transcript (*d*_*xy*_= Number of fixed SNPs / total number of SNPs). Then, CDS sequences were predicted using the TransDecoder software (http://transdecoder.github.io) and evolutionary rates were estimated with the KaKsCalculator program (Zhang et al., 2006). The estimated evolutionary rate between species was further used to investigate its relationship with the expression inheritance and regulatory divergence data.

### Molecular divergence and regulatory dynamics

To investigate potential associations between different categories of expression and molecular divergence, we implemented a GLM framework comparing variables of interest among categories/profiles. A comparative matrix containing all transcripts, including ED, inheritance modes, regulatory divergence categories, *cis* index of gene regulation (proportion of gene expression relative to *trans* effect for each transcript), and Ka/Ks estimates between species was generated. GLM analyses were performed using categories of regulatory divergence, inheritance modes, and tissues as independent variables, while both the number of transcripts detected, and the expression divergence were considered dependent variables. Then, the response variables were compared across categories of inheritance, reciprocal hybrids (OF and FO), and reproductive tissues. Because the number of transcripts detected under each category depends on the transcriptome length (number of assembled transcripts), we normalized all transcript counts to the number of assembled contigs per library. Since several variables are calculated as proportions (e.g., *cis* index, Ka/Ks, normalized number of transcripts), GLM analyses were performed after square-root transformation.

### Transcriptional annotation and functional analysis

Assembled transcriptomes were annotated using the Trinotate framework (http://trinotate.github.io). For this, we used predicted CDS regions from TransDecoder to perform homology searches against known sequence databases (*e.g.,* SWISSPROT) using the programs BLASTX (transcripts) and BLASTP (predicted CDS) (Camacho et al., 2009). HMMER software was used to identify conserved functional domains (Wheeler and Eddy, 2013) against the PFAM-A database. rRNA sequences were identified in transcripts using the RNAMMER tool (Lagesen et al., 2007). Finally, the signalP software (Almagro Armenteros et al., 2019) was used to identify signal peptides (secretion signals), while the tmHMM tool was used to identify contigs by predicting transmembrane domains (Krogh et al., 2001). All these searches were integrated into a Trinotate SQLite database to produce the final annotation file. Overrepresentation of specific categories of biological functions was investigated for genes following inheritance modes of gene expression and regulatory divergence using the GOseq R package framework (Young et al., 2010).

## Results

### Sequencing and trimming de novo assembly

Nearly 300 million paired-end read sequences were obtained from the Illumina runs, ranging from 11 to 24 million raw paired-end reads for each sample (Table S1). After trimming and filtering of sequence reads containing fixed SNPs between species, an average of 4 × 10^6^ unambiguously mapped reads were obtained for each library. Transcriptomes assembled *de novo* and corrected for redundancy and chimerism contained between 80 to 100 thousand transcripts per reproductive library. From these assembled transcripts, after final filtering of 3 cpm in at least two samples from each profile, we detected at least one fixed SNP between the parental species in between 9,709 to 10,628 transcripts. On average, we found 4.4 fixed SNPs per transcript between the parental species. When these SNPs were used to filter the read-count matrix used for ASE estimations, the proportion of filtered reads was slightly higher for *A. fraterculus* than *A. obliqua*. Moreover, hybrids between these species exhibited a nearly 1:1 ratio of mapping reads based on their parental origins (Table S1), as expected in absence of mapping bias. We then confirmed these expectations by simulating reads extracted from both assemblies (*D. fraterculus* and *D. obliqua* derived references) in the “diploid *Anastrepha* genome”. The same number of reads was simulated for each polymorphic site and then mapped back to the reference. As expected in the absence of mapping bias, transcripts in all samples followed an expression ratio of one.

### Expression divergence between A. fraterculus and A. obliqua

The number of transcripts with significant expression divergence (ED) between species was substantially different between reproductive transcriptomes of males and females (Figure S1a, Table S2 and S3): 2248 female reproductive transcripts (11%) exhibited ED between species, while 1177 ED genes (23%) were detected in males (Figure S1a, Table S2 and S3). These differences remained after accounting for library size (Table S3).

### ASE in hybrids

10 to 14% of filtered transcripts showed ASE in hybrids (Figure S1b, Table S2). Between 40 and 52% of these transcripts showed higher expression for *A. fraterculus* alleles in hybrid reproductive tissues (Figure S1b, Table S2). OF and FO hybrid females had similar relative expression distributions and differed from that of FO males (Figure S1b, Table S2). These differences between reciprocal crosses were, however, not observed when comparing the inheritance mode, or regulatory divergence of their gene expression (see below).

### Modes of expression inheritance

We classified transcripts according to their inheritance modes, based on pairwise comparisons between parents and hybrids (Figure 2a, S2). Hybrid expression was assessed using all reads per transcript, independent of their allelic origin. The expression of 84 to 94% of transcripts was conserved (Table S2). We used a GLM analysis to investigate differences in the distribution of the remaining transcripts across categories between male and female reproductive tissues (Table S3). After controlling for the total number of transcripts per library, the relative number of transcripts detected and the pairwise interactions between these factors were significantly different across categories and tissues (Table S3). The proportion of transcripts was substantially different across categories (Table S3). The majority of transcripts show dominance (more often of *A. fraterculus* over *A. obliqua*), followed by transgressive categories, with *overdominance* being significantly more common than *underdominance* (Figure 2a, S3). Pairwise interaction effects indicated that the number of transcripts, or their distribution across categories, is different between females and males (Figure 2a, S2). The number of detected transcripts was greater for females than males across categories (Figure 2a, S3). The direction of the cross (*e.g.*, FO or OF) did not significantly affect the number of transcripts detected under these categories.

**Figure 2.**
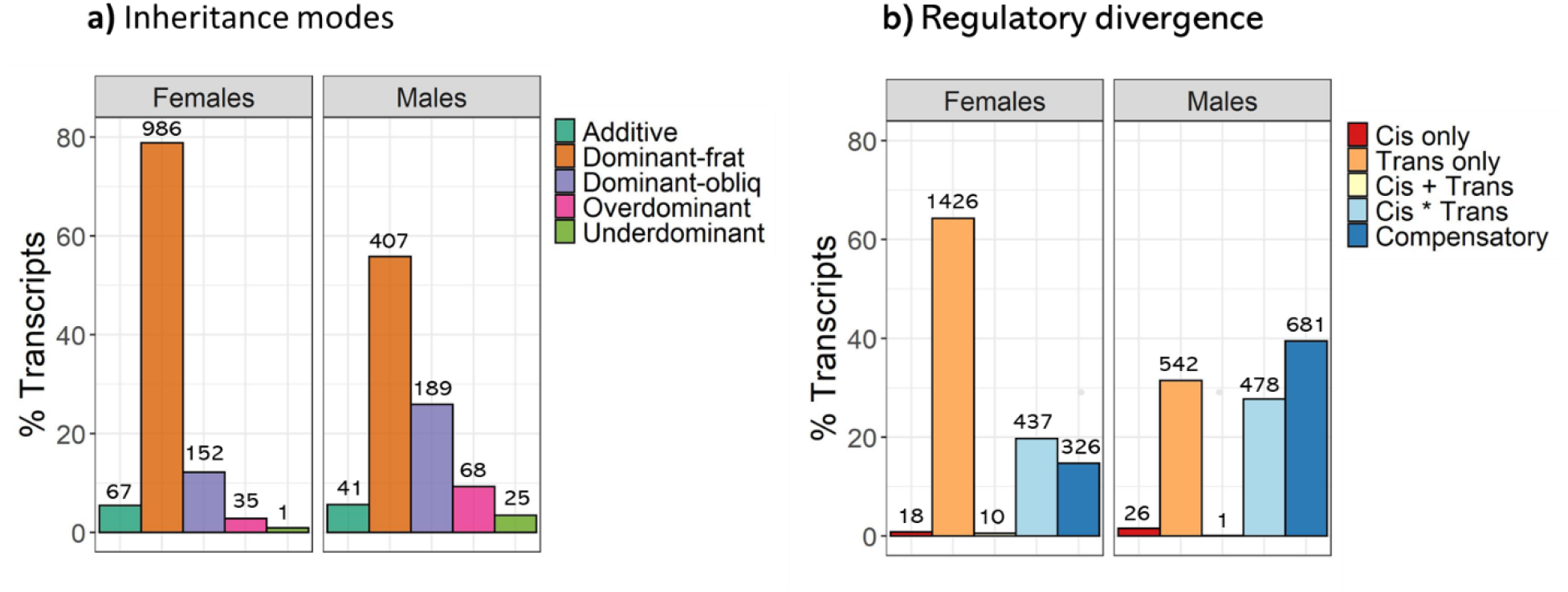
Classification of transcripts according to their a) mode of gene expression inheritance and b) mechanisms of regulatory divergence. Barplots show the distribution of ED genes for each category separated by sex, after excluding conserved and ambiguous transcripts (in the case of regulatory categories). These categories are based on transcript showing significant differential expression between the parental species. The number of transcripts in each category is indicated on top of each bar.

### Regulatory divergence inferred from ASE

Patterns of ASE in hybrids and gene expression in the parental species were used to infer *cis* and *trans*-regulatory effects for each transcript (Figure 2b, S3). To unambiguously assign alleles to parental species, only transcripts with expression in both parents were used. 55 to 61% of transcripts were conserved (Table S2), showing no expression divergence between species and no significant ASE in hybrids, while about 22% of transcripts were classified as ambiguous (Table S2). The remaining transcripts were classified under five regulatory categories whose distribution across profiles was compared using a GLM analysis (Table S3). After controlling for the total number of transcripts, the relative number of transcripts detected was significantly influenced by categories and reproductive tissues (males and females) as well as their pairwise interactions (Table S3). The number of transcripts was substantially different across categories, with the majority of transcripts for *trans*-only and interactions between *cis* and *trans* (*cis * trans* or *compensatory*), while only a few transcripts showed evidence for *cis*-only or *cis + trans*-regulatory effects (Figure 2b, S3). Pairwise interactions indicate that the total number of transcripts or their distribution across categories is different between males and female reproductive tissues (Figure 2b, S3). The number of *trans-*only transcripts was greater on female than on male reproductive tissues. On the other hand, the number of transcripts with evidence for *cis* and *trans* interactions (*cis* * *trans* or *compensatory*) was higher for male profiles (Figure 2b, S3). The direction of the cross (*e.g.*, FO or OF) did not significantly affect the number of transcripts detected under these categories.

### Expression divergence across categories of inheritance and regulatory divergence

Gene expression divergence (ED) was substantial across categories of inheritance modes (Figure 3a) and regulatory mechanisms (Figure 3b) between species (Table S3). ED tended to be higher for *additive* transcripts, followed by dominant and finally transgressive transcripts (over and underdominant) (Figure 3a). No significant differences between regulatory categories were detected between male and female reproductive transcriptomes, while GLM showed evidence for pairwise interactions between them (Table S3). Interactions showed that ED was substantially higher for transcripts exhibiting *cis* + *trans*regulatory divergence, while the lowest ED was detected for transcripts in the *compensatory* category and no substantial differences across the remaining mechanisms (Figure 3b). The direction of the cross (*e.g.*, FO or OF) did not significantly affect the level expression divergence detected under these categories.

**Figure 3.**
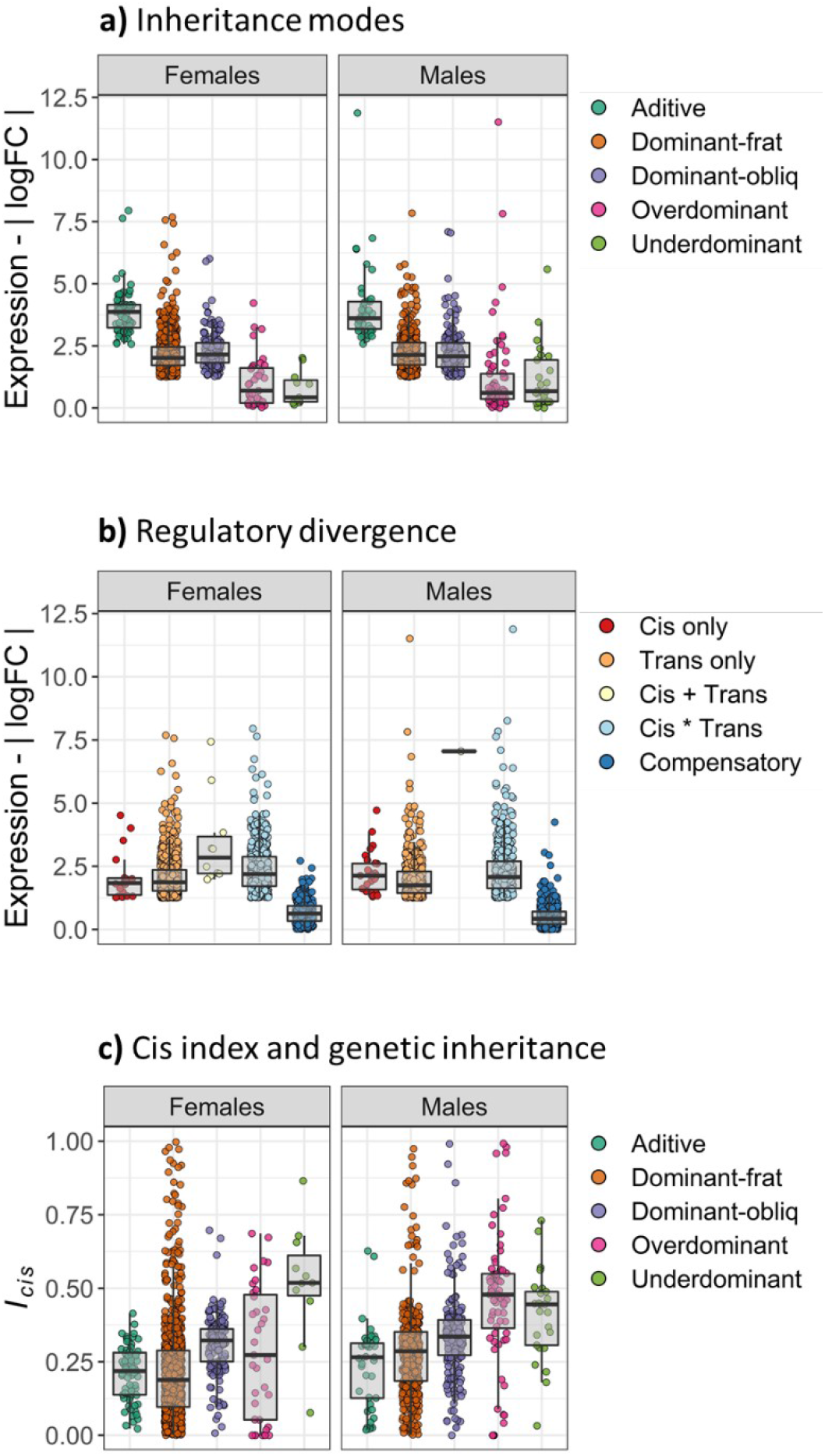
Patterns of expression divergence across categories of genetic inheritance and gene regulation between *A. fraterculus* and *A. obliqua*. ED between the species was compared across **a)** modes of expression inheritance and, **b)** regulatory divergence. Boxplots compare variation of absolute relative expression (a and b) between parental species (| log_2_FC *frat* / *obliq* |). **c)** The *cis* index *(I*_*cis*_) of regulatory divergence was compared across categories of genetic inheritance in hybrids. Boxplots compare the average variation of *cis* index per transcript.

### Cis index and inheritance modes

We compared the *cis* index of gene regulation (relative size of the *cis* to *trans* effect per transcript) across inheritance modes of expression (Table S4). The GLM results showed that the *cis* index was significantly different across categories of inheritance modes and reproductive tissues, as well as their interactions (Table S4, Figure 3c). The *cis* index tended to be higher for transgressive transcripts (over and underdominant), followed by dominant transcripts (often higher in dominant-obliq than dominant-frat), and finally additive transcripts (Figure 3c). The direction of the cross (*e.g.*, FO or OF) did not significantly affect the cis index detected across categories of expression inheritance.

### Relationship between molecular divergence, expression divergence, and allele imbalance in hybrids

The level of molecular divergence as obtained from Ka, Ks and its ratio (Ka/Ks), followed similar patterns, being significantly higher for genes with expression divergence (ED) between the species when compared to *conserved* genes (Figure 4a). The molecular divergence estimates for these three indices differed between male and female reproductive transcriptomes, with males exhibiting significantly higher values when compared to those of females. Similarly, molecular divergence estimates were consistently higher for genes with significant allele imbalance in hybrids (AI), but no significant differences between males and females were found (Figure 4b).

**Figure 4.**
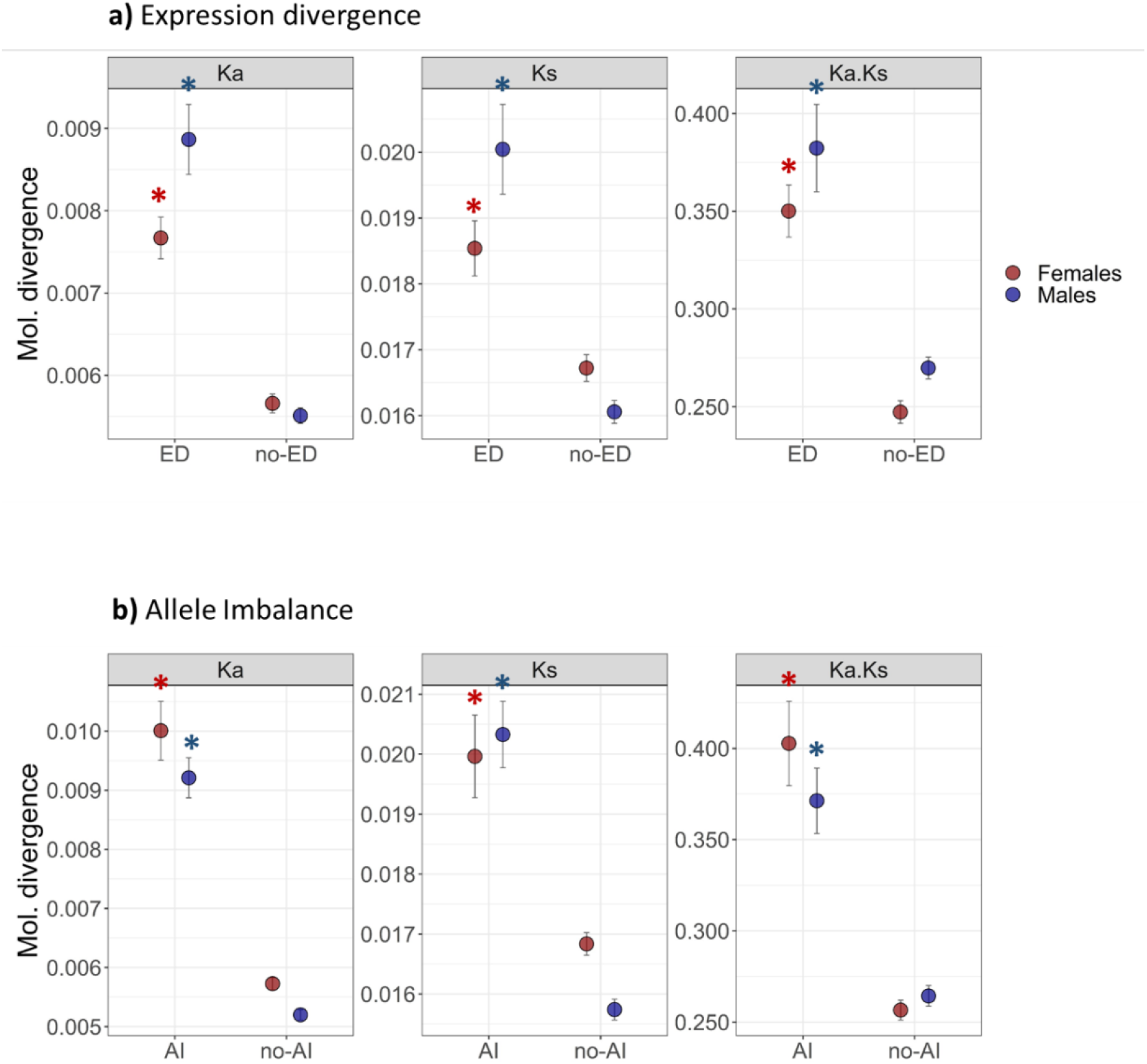
Patterns of molecular divergence associated with expression divergence and allele imbalance in hybrids between *A. fraterculus* and *A. obliqua*. The plots compare the average molecular divergence (Ka, Ks and Ka/Ks) for transcripts with significant **a)** expression divergence (ED), and **b)** allele imbalance (AI). Significant comparisons following GLM analysis between groups are indicated for males and females are indicated with *.

### Relationship between molecular divergence and expression inheritance in hybrids

When comparing molecular divergence across categories of expression inheritance, we found that Ka and Ks estimates followed slightly different patterns. All categories of inheritance show significantly higher Ka divergence when compared to *conserved* genes, and additive genes show the highest levels of molecular divergence. Likewise, dominant and transgressive genes showed significantly higher Ks divergence estimates, whereas *additive* genes are not significantly different from *conserved* genes. All categories then have higher Ka/Ks ratios when compared to conserved genes, although male additive genes and female transgressive genes values are not significant, perhaps due to a large variance. Interestingly, *transgressive* genes show the highest molecular divergence for Ks values in males and females. (Figure 5a).

**Figure 5.**
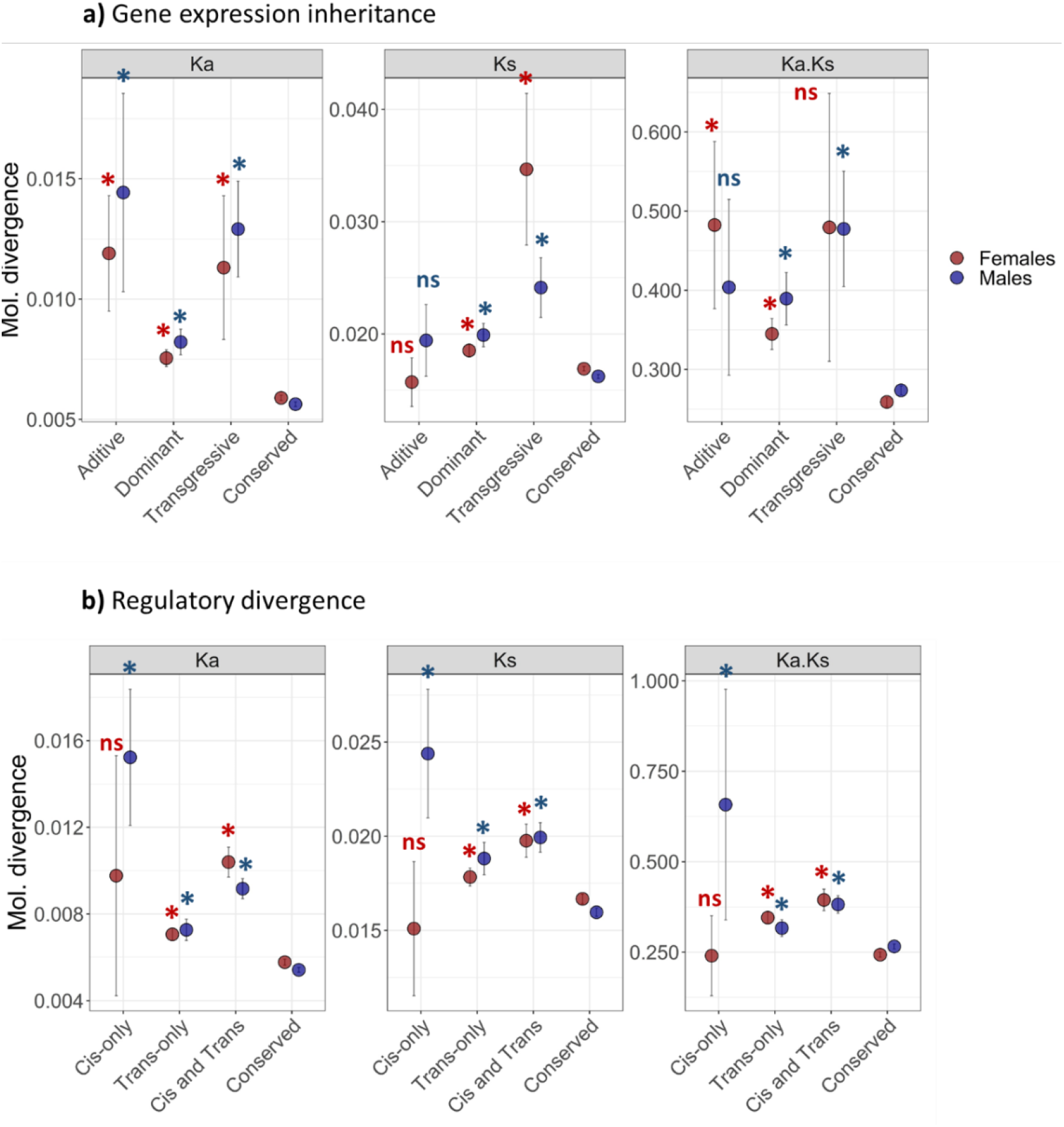
Patterns of molecular divergence associated with expression inheritance and regulatory divergence between *A. fraterculus* and *A. obliqua*. The plots compare the average molecular divergence (Ka, Ks and Ka/Ks) across different classes of **a)** gene expression inheritance, and **b)** regulatory divergence. Significant comparisons following GLM analysis for the comparison between each class and the conserved group of genes for males and females are indicated with *.

### Relationship between molecular and regulatory divergences

Estimates of molecular divergence are also slightly different for categories of regulatory divergence, showing consistent patterns across Ka, Ks and Ka/Ks ratios (Figure 5b). In this case, all regulatory divergence categories show significantly higher molecular divergence when compared to conserved genes. The only exception is that female cis-only genes are not significantly different to conserved genes, whereas male cis-only genes exhibit the highest molecular divergence for *cis*-only genes (Figure 5b), even higher than that of *transgressive* genes (Figure 5a). These results suggest that transcriptional differences in expression and regulation between *Anastrepha* species explain patterns of molecular divergence.

### Functional analysis

To analyze the functional pathways associated with gene expression inheritance and regulatory divergence between *A. fraterculus* and *A. obliqua*, we performed gene ontology (GO) enrichment analysis (Figure 6). We found evidence for functional specialization played by genes with *trans*-only divergence and *dominant-frat* expression in hybrids between these *Anastrepha* species (Figure 6a). While the rest of the categories did not show significant gene enrichments in female transcriptomes, these two sets of genes were strongly associated with important functions previously linked to the female reproductive tract of numerous insect species (Figure 6b). These transcriptional changes are enriched by the following three groups of functional pathways (Figure 6b): i) epithelial modifications, ii) protease/protease inhibitors, and iii) immune/defense response. Similarly, male transcriptomes were significantly enriched by at least three functional pathways that have been previously associated with male reproductive tissues.

**Figure 6.**
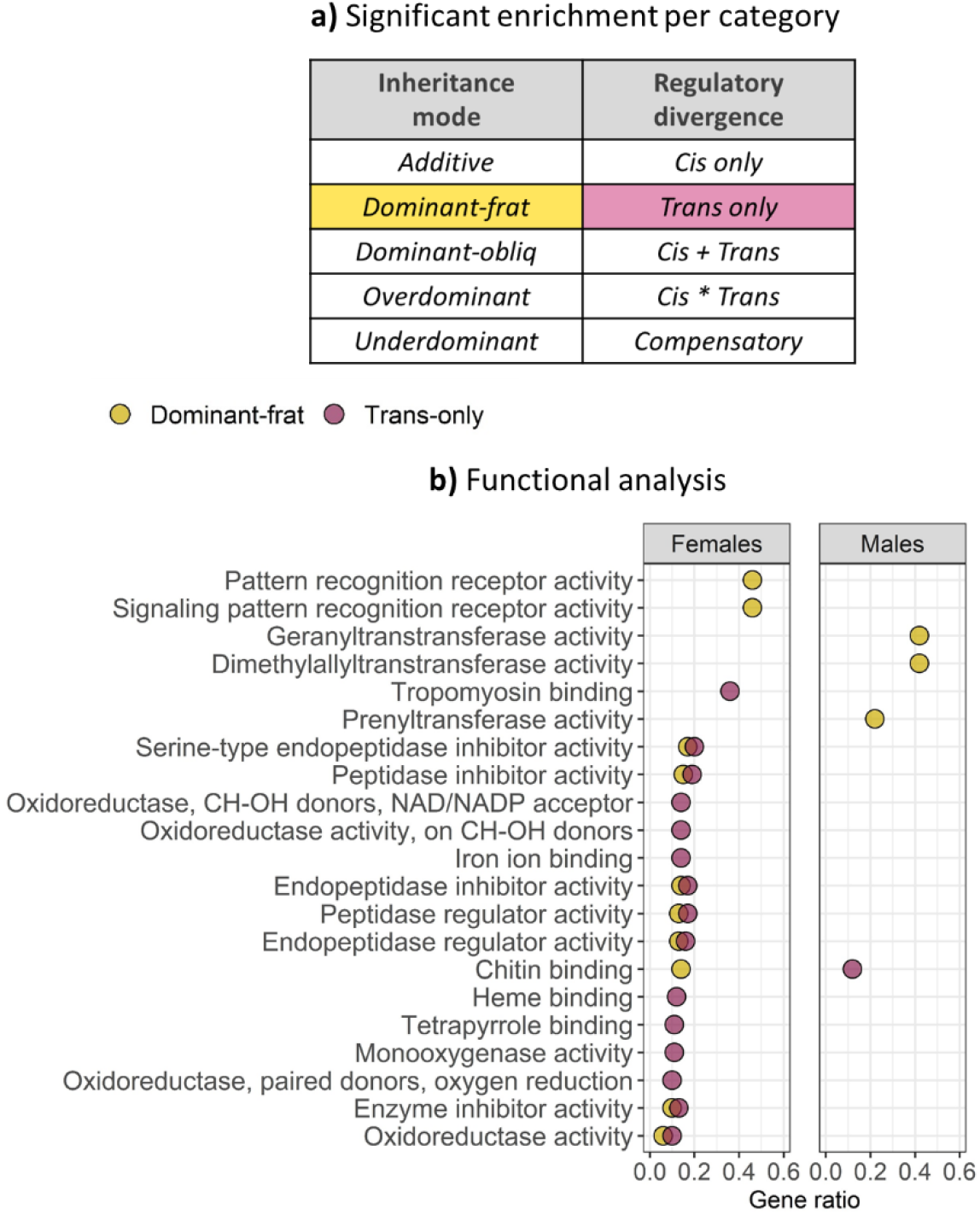
Functional analysis of male and female reproductive transcripts following modes of expression inheritance and regulatory divergence between *A. fraterculus* and *A. obliqua*. **a)** Gene ontology enrichments were significant for genes with *dominant-frat* expression in hybrids as well as those with *trans*-only regulatory divergence. **b)** Functional analysis indicates that these categories are enriched for regulatory networks associated with male and female sexual interactions in insects. Gene ratio of significantly detected genes within each enriched category is indicated (Gene ratio = significant genes in category / total number of genes in category). All significant comparisons with following *GLM* analysis are indicated with *.

## Discussion

Using patterns of transcriptional divergence and allele-specific expression (ASE) in F_1_ hybrids between species connected by gene flow (*A. fraterculus* and *A. obliqua*) we have been able to characterize regulatory divergence and expression inheritance, and link these to sequence divergence. Expression inheritance characterizes the relationship between the pattern of gene expression in hybrids and that of the parental species, while regulatory divergence uses the pattern of ASE in hybrids to characterize the expression regulation of the gene copies coming from the two parental species. We find that transcriptional patterns in the hybrids show a mosaic between those typically observed within and between allopatric species. The majority of differentially expressed genes between the species showed an interaction between *cis* and *trans* regulatory divergence, particularly a *compensatory* pattern, which is more frequently seen in interspecific hybrids (Wittkopp et al., 2008; Coolon et al., 2014). There is also a relatively high proportion of genes that show *trans*-only regulatory divergence and *dominance* expression inheritance, which is consistent with the pattern seen in within-species ASE studies (Wittkopp et al., 2008; Suvorov et al., 2013).

### Gene-expression inheritance in hybrids

We detected all possible modes of expression inheritance, with a similar distribution across male and female reproductive transcriptomes. Most transcripts that are differentially expressed fall into one of the *dominant* categories (80-95%), followed by the *transgressive* (*over-* or *underdominant*) modes and *additive* mode of inheritance (2-6%), and that is true for both sexes. Our study did not find *transgressive* transcripts to be as common as described in other interspecific hybrids (Landry et al. 2007; Wittkopp et al. 2008b). Indeed, the proportion of *transgressive* transcripts (3-13%) is over three times lower than that observed in interspecific *Drosophila* hybrids (35-69%) (Ranz et al., 2004; McManus et al., 2010), and even lower than some intraspecific crosses (Bell et al., 2013). Transgressive phenotypes are often assumed to be a consequence of *transgressive* expression (Rieseberg et al. 2003; Landry et al. 2005; Swanson-Wagner et al. 2006), but the pattern of expression divergence we see across inheritance categories does not support this hypothesis. Rather, we see non-additively inherited transcripts exhibiting much lower expression divergence than *additive* transcripts. Our results suggest that genes with *transgressive* expression are more likely to show low expression divergence while additively inherited transcripts show the highest expression divergence. Very few genes show *transgressive* expression in hybrids when compared to non-transgressively inherited transcripts, which may be consistent with the reduced genetic divergence between *A. fraterculus* and *A. obliqua* and the fact that lab generated hybrids are viable, fertile, and show no visible abnormalities aside from the Haldane’s rule that characterizes the inviability of the male sex in one of the interspecific crosses (Selivon et al., 1999; Santos et al., 2001; Rull et al., 2017). More distantly related species, such as *C. briggsae* and *C. nigoni*, have a high proportion of transgressively expressed genes, as well as *cis-trans* compensatory changes (Sanchez-Ramirez et al., 2021).

### Regulatory divergence

Comparisons across *Drosophila* species (Coolon et al., 2014; Meiklejohn et al., 2014) have shown that the proportion of genes showing *cis* or *trans* regulation can change with the level of species divergence. Since divergence in *cis* regulation is expected to result from accumulated changes at individual transcripts that are independent of background (Coolon et al., 2014; Signor and Nuzhdin, 2018), these changes may require more time to accumulate. The contribution of *trans* regulation to divergence likely depends on whether a given gene is *trans* regulated or is the source of *trans* regulation. *Trans* regulators are likely to be constrained by their pleiotropic effects given that their influence can potentially cascade across multiple target genes (Wittkopp et al., 2008; Mack and Nachman, 2017), while genes that are regulated in *trans* may not show this same constraint.

The majority of differentially expressed transcripts shows *cis*-*trans* interactions (*i.e.*, *cis * trans* or *compensatory*), with opposing effects on gene expression with a large proportion of transcripts showing compensatory regulation in hybrids. This pattern, along with the high proportion of genes showing conserved expression, provides strong evidence for evolutionary constraint on gene expression (Bell et al., 2013; Chen et al., 2015; Graze et al., 2012; Meiklejohn et al., 2014; Ortiz-Barrientos et al., 2007; Signor and Nuzhdin, 2018) in a substantial number of genes of both male and female reproductive transcriptomes. This general pattern is widespread, being also found in yeast, worms, *Drosophila,* and mice (Mack and Nachman, 2017; Signor and Nuzhdin, 2018). It has been explained by coadaptation of regulatory elements mediated by stabilizing selection of gene expression. Despite the extensive evidence for this pattern, its evolutionary significance and the relative contribution of selection or parallel evolution are usually unexplored (Signor and Nuzhdin, 2018). So far, available models and theoretical considerations suggest that independent evolution of *trans* elements evolves following speciation (Ortiz-Barrientos et al., 2007; Signor and Nuzhdin, 2018). Then, stabilizing selection on gene expression favors compensation mediated by *cis-* elements of target genes (Ortiz-Barrientos et al., 2007; Fear et al., 2016; Mack and Nachman, 2017; Signor and Nuzhdin, 2018).

### Transcriptional incompatibilities and molecular divergence

We found significant association between rates of molecular evolution of individual genes and their level of expression divergence between species, as well as with the extent of allelic imbalance in hybrids. These results support the hypothesis that protein sequence and expression divergence are influenced by similar selective processes (Go and Civetta, 2020). Within these genes, we discovered that genes associated with *transgressive* expression and those showing *cis*-only regulatory divergence exhibit the greatest molecular divergence. Therefore, as expected from previous findings and models of hybrid incompatibility (Coyne and Orr, 1997; Landry et al., 2005; Ortiz-Barrientos et al., 2007; Presgraves, 2008; Maheshwari and Barbash, 2011; Masly and Presgraves, 2012; Victoria Cattani and Presgraves, 2012; Turelli et al., 2014; Lopez-Maestre et al., 2017), our results suggest that the underlying causes of genetic incompatibility between *Anastrepha* species arise primarily from positive selection or relaxation of selective constrains in both *transgressive* and *cis-*regulated genes. More interestingly, these groups of genes also show the highest difference in patterns of molecular evolution between sexes, with males showing higher evolutionary rates than females. Moreover, we found evidence of selection for male *transgressive* and *cis*-regulated genes, suggesting positive selection or relaxed constraints driving resistance to gene flow in male but not in female reproductive transcriptomes.

Female *transgressive* genes exhibit the highest level of divergence at synonymous sites, suggesting more resistance to gene flow. These transcripts also show elevated divergence at nonsynonymous sites, but do not show an elevated relative rate of nonsynonymous change (i.e., their Ka/Ks ratio is not significantly elevated over that of *conserved* transcripts). This suggests that the nonsynonymous changes lead to transcriptional incompatibilities in female hybrids, but they are not experiencing recurrent divergent positive selection. Likewise, male *transgressive* genes also show *K*a and *Ks* levels significantly higher than that of *conserved* genes, but in this case, their ratio (*Ka/Ks*) is also significantly higher, indicating a more relevant role of positive or relaxed selection. *cis*-regulated male genes show the highest molecular divergence for synonymous and nonsynonymous changes, leading to the highest Ka/Ks estimates, whereas *cis*-regulated female genes show higher *Ka* and *Ks*, but not Ka/Ks, as it was seen for *transgressive* female genes. This particular result suggests one possible explanation for the Haldane’s rule detected in these species, where hybrids involving *A. obliqua* females (Selivon et al., 1999; Santos et al., 2001; Rull et al., 2017) do not produce male offspring due to a preponderance of *cis-*only regulation or *transgressive* expression in male hybrids, which are presumably major causes of genetic incompatibility between these species.

Patterns of molecular evolution in *transgressive* and *cis*-regulated transcripts are consistent with the fact that these two groups of genes are associated with hybrid misexpression and are likely to influence hybrid fitness. However, they also represent the smallest fraction of transcriptional differentiation between *Anastrepha* species. While most of the transcriptional landscape is dominated by genes that do not show differences between the species, those that differ, tend to show a pattern of *compensatory* regulation and *dominant* inheritance in hybrids. This is expected for species in isolation with gene flow, since their hybrids must overcome their transcriptional incompatibilities to account for the presence of introgression between the species (Díaz et al., 2018). The larger proportion of *trans*-only inheritance was found on female genes, whereas male genes have a higher proportion of compensatory changes, could reflect lower rates of evolution on female genes (Goncalves et al., 2012). Higher rates of evolution on male genes, especially on spermatogenesis genes that are more prone to experiencing compensatory divergence for X-linked genes (Sanchez-Ramirez et al., 2021), may underlie the higher proportion of compensatory changes on male genes as well as the male hybrid expression disfunction behind Haldane’s rule.

The difference in genetic constraint between *cis* and *trans* regulation suggests that these mechanisms might not only play different roles in adaptation and the evolution of genetic incompatibilities (Ortiz-Barrientos et al., 2007; Mack and Nachman, 2017; Signor and Nuzhdin, 2018), but also might be differentially affected by gene flow. For example, numerous studies have suggested that *cis* effects are often more associated with higher molecular and expression divergences when compared to *trans* effects (Wittkopp et al., 2004, 2008; Meiklejohn et al., 2014; Fear et al., 2016). Even though our results indicate that genes with significant expression divergence tend to show higher molecular divergence on both sexes, only on males we observe that *cis*-only regulated genes show higher evolutionary rates than *trans*-regulated genes. However, expression divergence between *A. fraterculus* and *A. obliqua* seems better predicted by the interactions between *cis* and *trans* elements. For example, *compensatory* transcripts do not exhibit expression divergence, while *cis* + *trans* interactions exhibit the most dramatic changes in expression divergence.

Despite substantial gene flow previously reported between these species in both directions (Diaz et al., 2016; Scally et al., 2016; Congrains et al., 2021), we detected a preponderance of *dominance* in the expression of *A. fraterculus* genes over *A. obliqua* genes. Functional analyses on these genes found *dominant-frat* and *trans*-only genes enriched for regulatory networks that have been associated multiple times with the female postmating response in insects (Mack et al., 2006; Kocher et al., 2008; Alfonso-Parra et al., 2016; Al-Wathiqui et al., 2016; Thailayil et al., 2018; Fowler et al., 2019; Liu and Hao, 2019; Gao et al., 2020). Similar studies on males detected that these same categories were enriched for regulatory networks associated with spermatogenesis (Dugan and Allen, 1995; Adolphsen et al., 2012). These results clearly show that female (*e.g.*, proteases/inhibitors or immune/defense functions) and male genes (*e.g.*, Prenyl- Dimethylallyl- and Geranyl-transferases) associated with reproduction are likely part of the “genomic islands of speciation” in species undergoing gene flow but are not as divergent and potentially incompatible as *cis* (in males) *or cis-trans* interacting genes (females).

The expression of reproductive genes in hybrids are essential to modulate introgression (Signor and Nuzhdin, 2018). Models of postzygotic incompatibilities predict that reproductive genes are affected by the genetic incompatibilities when combining the genomes of the two parental species in one individual (Coyne and Orr, 1997; Landry et al., 2005; Ortiz-Barrientos et al., 2007; Presgraves, 2008; Maheshwari and Barbash, 2011; Masly and Presgraves, 2012; Victoria Cattani and Presgraves, 2012; Turelli et al., 2014; Lopez-Maestre et al., 2017). We have demonstrated that reproductive genes are in fact associated with transcriptional differentiation having the lesser impact on hybrid fitness. Differentially expressed reproductive genes are significantly more associated with *dominance* of *A. fraterculus*-like expression and *trans*-regulation in hybrids, showing significant enrichment for gene ontology categories previously associated with female (Mack et al., 2006; Kocher et al., 2008; Thailayil et al., 2018; Fowler et al., 2019; Liu and Hao, 2019; Gao et al., 2020) and male reproductive pathways (Dugan and Allen, 1995; Adolphsen et al., 2012). This suggests that these regulatory mechanisms are competitively stronger than those in *A. obliqua* when they are together in the same hybrid. Other studies in hybrids have shown biased expression of one genome over another (Yoo et al., 2013; Banho et al., 2021). Not only does *A. fraterculus* appear to be a more variable species when compared to *A. obliqua* (Solferini and Morgante, 1987; Rezende Borges et al., 2016), but it has also been reported as a more variable lineage, to the point that it is currently considered a species complex in itself, with several entities detected in South America (Hernández-Ortiz et al., 2004, 2015; Hendrichs et al., 2015; Vaníčková et al., 2015). This might indicate that observed *trans*-regulatory interactions have diverged in *A. fraterculus* with respect to *A. obliqua* and might be an important characteristic of their rapid radiation, particularly for reproductive tissues expressing rapid evolving genes. More interestingly, these results point to *trans*-only regulation as one of the mechanisms by which species can evolve in the presence of gene flow and divergent selection, where the regulatory mechanisms of one species takes control of reproductive transcriptional networks, and therefore buffering hybrid misexpression to cause hybrid or reproductive breakdown.

## Acknowledgments

We would like to thank FAPESP (Fundaçao do Amparo à Pesquisa do Estado de São Paulo) Grant #2010/20455-4 and the Science without Borders program at CAPES (Processo PVE 056/2013) for financial support. We would also like to thank Drs. Roberto Zucchi and Keiko Uramoto, from University of São Paulo, for help in identifying several of specimens here used.

