## Supplementary Material for "Patterns of regulatory divergence and gene expression in hybrids are associated with molecular evolution in species undergoing gene flow"

### TABLES

**Table S1.** Sample profiles used for RNA-Seq analysis. Each replicate corresponds to a pool of five individuals.

| ***Sample*** | | ***Replicates*** | ***Mean read pairs*** | ***After trimming*** | ***% aligned*** |
|---|---|---|---|---|---|
| ***A. frateculus*** | | | | | |
| *Head* | ♀ | 3 | 14267520 ± 155539 | 12663188 ± 401935 | 80.3 |
| | ♂ | 3 | 13376642 ± 128254 | 11578222 ± 99084 | 79.8 |
| *Reproductive* | ♀ | 3 | 13883827 ± 319748 | 11944386 ± 292906 | 83.3 |
| | ♂ | 3 | 13249470 ± 719148 | 11470063 ± 632289 | 80.9 |
| ***A. obliqua*** | | | | | |
| *Head* | ♀ | 3 | 11915997 ± 502234 | 10530475 ± 560495 | 80.2 |
| | ♂ | 3 | 13726190 ± 600766 | 12349894 ± 530269 | 83.4 |
| *Reproductive* | ♀ | 3 | 13132508 ± 1307881 | 11770098 ± 1253708 | 84.3 |
| | ♂ | 3 | 12357116 ± 296177 | 11140920 ± 301655 | 84.5 |
| *Hybrid: ♂ frat x ♀ obliq* | | | | | |
| *Head* | ♀ *(only)* | 3 | 22351663 ± 1979872 | 18989084 ± 1767636 | 70.9 |
| *Reproductive* | ♀ *(only)* | 3 | 27316867 ± 4083976 | 22987040 ± 3194692 | 73.7 |
| *Hybrid: ♀ frat x ♂ obliq* | | | | | |
| *Head* | ♀ | 3 | 13222834 ± 662739 | 10964238 ± 666362 | 71.0 |
| | ♂ | 3 | 13943759 ± 685806 | 11690966 ± 552469 | 70.7 |
| *Reproductive* | ♀ | 3 | 12418845 ± 464623 | 104923612 ± 499870 | 74.9 |
| | ♂ | 3 | 13713823 ± 709307 | 11529501 ± 585536 | 72.1 |

**Table S2.** Summary of differentially expressed transcripts between parental species, allelic imbalance and their classifications according to inheritance modes and regulatory mechanisms responsible for divergence.

| **Tissue** | *Head* | *Head* | *Head* | *Reprod.* | *Reprod.* | *Reprod.* |
|---|---|---|---|---|---|---|
| **Sex** | *Female* | *Female* | *Male* | *Female* | *Female* | *Male* |
| **Cross** | ♂ *f* * ♀ *o* | ♀ *f* * ♂ *o* | ♀ *f* * ♂ *o* | ♂ *f* * ♀ *o* | ♀ *f* * ♂ *o* | ♀ *f* * ♂ *o* |
| *Total contigs* | 14627 | 14627 | 15066 | 9709 | 9709 | 10628 |
| ***Expression divergence*** | | | | | | |
| FDR $\alpha = 0.05$ | 2869 | 2869 | 533 | 3513 | 3513 | 2360 |
| *frat* parent > *obliq* parent | 1649 | 1649 | 350 | 2076 | 2076 | 1143 |
| FDR $\alpha = 0.05$ and logFC > 1.25 | 1552 | 1552 | 533 | 2248 | 2248 | 1177 |
| *frat* parent > *obliq* parent | 940 | 940 | 350 | 1540 | 1540 | 645 |
| ***ASE*** | | | | | | |
| FDR $\alpha = 0.05$ | 2496 | 1663 | 2254 | 1919 | 1560 | 1765 |
| *frat* allele > *obliq* allele | 1314 | 803 | 1056 | 1092 | 846 | 984 |
| FDR $\alpha = 0.05$ and logFC > 1.25 | 1852 | 1232 | 1980 | 1113 | 943 | 1437 |
| *frat* allele > *obliq* allele | 1035 | 594 | 907 | 685 | 535 | 804 |
| ***Inheritance modes*** | | | | | | |
| Additive | 34 | 35 | 7 | 30 | 67 | 41 |
| Dominant-frat | 494 | 463 | 46 | 1330 | 986 | 407 |
| Dominant-obliq | 236 | 280 | 202 | 99 | 152 | 189 |
| Overdominant | 79 | 161 | 11 | 33 | 35 | 68 |
| Underdominant | 15 | 21 | 3 | 11 | 11 | 25 |
| Conserved | 13769 | 13667 | 14797 | 8206 | 8458 | 9898 |
| ***Regulatory divergence*** | | | | | | |
| *Cis*-only | 18 | 11 | 16 | 31 | 18 | 26 |
| *Trans*-only | 506 | 738 | 225 | 1300 | 1426 | 542 |
| *Cis + Trans* | 3 | 2 | 0 | 4 | 10 | 1 |
| *Cis * Trans* | 783 | 560 | 261 | 500 | 437 | 478 |
| Compensatory | 662 | 464 | 1478 | 381 | 326 | 681 |
| Conserved | 10036 | 10089 | 8883 | 5390 | 5321 | 6522 |
| Ambiguous | 2619 | 2763 | 4203 | 2103 | 2171 | 2378 |

**Table S3. Generalized linear model analysis of the number and expression of transcripts across categories of expression inheritance and regulatory divergence.** Both the number of transcripts and their expression were compared for each of the categories of inheritance mode and regulatory divergence. Models also included as the tissue type (head vs. reproductive) and sex. Data for number of transcripts were normalized across libraries and then square root transformed.

| *Effect* | *Number of transcripts* | | | | *Expression divergence* | | | |
|---|---|---|---|---|---|---|---|---|
| | *Df* | *Resid. Df* | *F* | *P* | *Df* | *Resid. Df* | *F* | *P* |
| ***Inheritance modes*** | | | | | | | | |
| Category | 4 | 25 | 122.3 | ***< 0.001*** | 4 | 5566 | 21.3 | ***< 0.001*** |
| Tissue | 1 | 24 | 38.5 | ***< 0.001*** | 1 | 5565 | 76.0 | ***< 0.001*** |
| Sex | 1 | 23 | 31.1 | ***< 0.001*** | 1 | 5564 | 34.9 | ***< 0.001*** |
| Category * Tissue | 4 | 19 | 26.4 | ***< 0.001*** | 4 | 5560 | 23.5 | ***< 0.001*** |
| Category * Sex | 4 | 15 | 16.9 | ***< 0.001*** | 4 | 5556 | 10.4 | ***< 0.001*** |
| Tissue * Sex | 1 | 14 | 5.3 | ***0.045*** | 1 | 5555 | 22.0 | ***< 0.001*** |
| Category * Tissue * Sex | 4 | 10 | 1.9 | 0.183 | 4 | 5551 | 1.0 | 0.429 |
| ***Regulatory divergence*** | | | | | | | | |
| Category | 4 | 25 | 280.1 | ***< 0.001*** | 4 | 11883 | 181.1 | ***< 0.001*** |
| Tissue | 1 | 24 | 40.8 | ***< 0.001*** | 1 | 11882 | 1.5 | 0.228 |
| Sex | 1 | 23 | 7.6 | ***0.021*** | 1 | 11881 | 0.8 | 0.375 |
| Category * Tissue | 4 | 19 | 25.7 | ***< 0.001*** | 4 | 11877 | 40.0 | ***< 0.001*** |
| Category * Sex | 4 | 15 | 29.8 | ***< 0.001*** | 4 | 11873 | 19.7 | ***< 0.001*** |
| Tissue * Sex | 1 | 14 | 0.9 | 0.371 | 1 | 11872 | 39.6 | ***< 0.001*** |
| Category * Tissue * Sex | 4 | 10 | 4.2 | ***0.029*** | 3 | 11869 | 27.0 | ***< 0.001*** |

Significant *P*-values are highlighted in bold

**Table S4. Generalized linear model analysis of the proportion of expression divergence due to *cis* effects relative to *trans* effects.** The *cis* index was estimated per transcript and compared across categories of inheritance modes, tissue and sex. Data for the *cis* component were normalized following square root transformation.

| *Effect* | *Df* | *Resid. Df* | *F* | *P* |
|---|---|---|---|---|
| Inheritance mode | 4 | 5566 | 190.7 | ***< 0.001*** |
| Tissue | 1 | 5565 | 137.0 | ***< 0.001*** |
| Sex | 1 | 5564 | 58.7 | ***< 0.001*** |
| Inheritance * Tissue | 4 | 5560 | 13.5 | ***< 0.001*** |
| Inheritance * Sex | 4 | 5556 | 7.9 | ***< 0.001*** |
| Tissue * Sex | 1 | 5555 | 20.5 | ***< 0.001*** |
| Inheritance * Tissue * Sex | 4 | 5551 | 8.9 | ***< 0.001*** |

Significant *P*-values are highlighted in bold

**FIGURES**

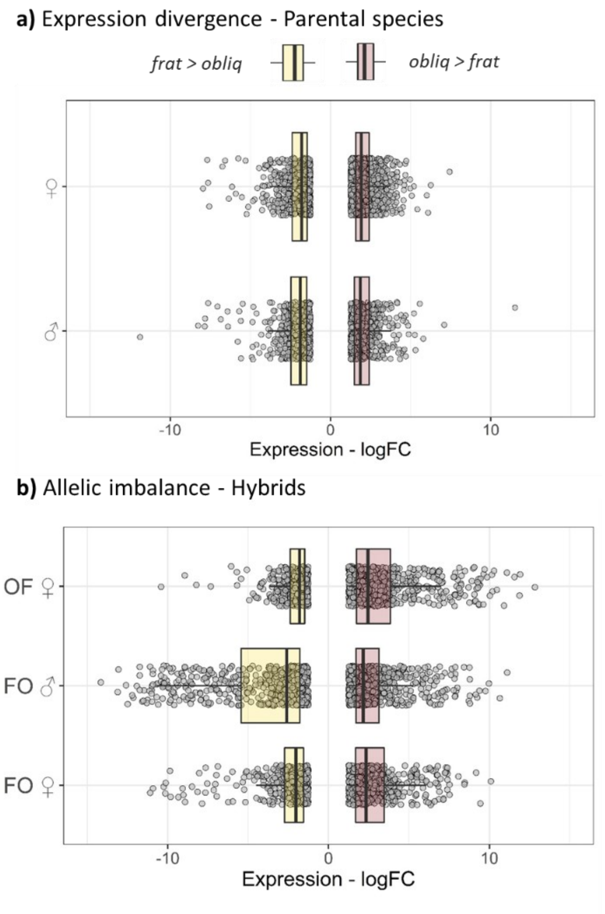


**Figure S1. Expression divergence and alllic imbalance in hybrids between between *A. fraterculus* and *A obliqua*. a)** Boxplots show the distribution of transcripts with expression divergenc (*$log_2FC$*) between the parental species for each reproductive tissue. **b)** Boxplots show distribution of transcripts with the allelic imbalance (*$log_2FC$*) between alleles in $F_1$ hyrbids for each direction of the cross (*e.g.*, FO vs OF, see methods).

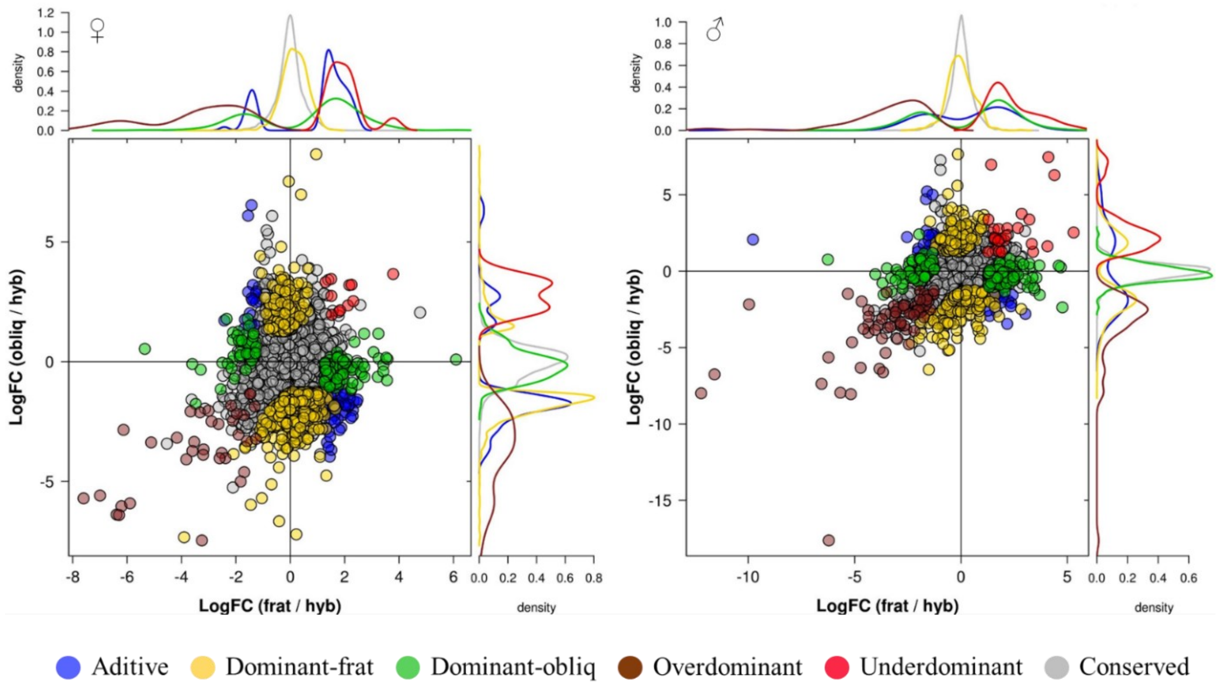


**Figure S2. Modes of expression inheritance in hybrids between *A. fraterculus* and *A. obliqua***. Scatterplots compare the relative expression of either parent (*A. fraterculus* / *A. obliqua*) and the hybrid [$log_2FC(frat/hyb)$ and $log_2FC(obliq/hyb)$], where hybrid expression is the sum of expression of the two alleles in hybrid (*hyb = hyb-frat + hyb-obliq*). These results were used to sort genes into categories based on their inheritance modes.

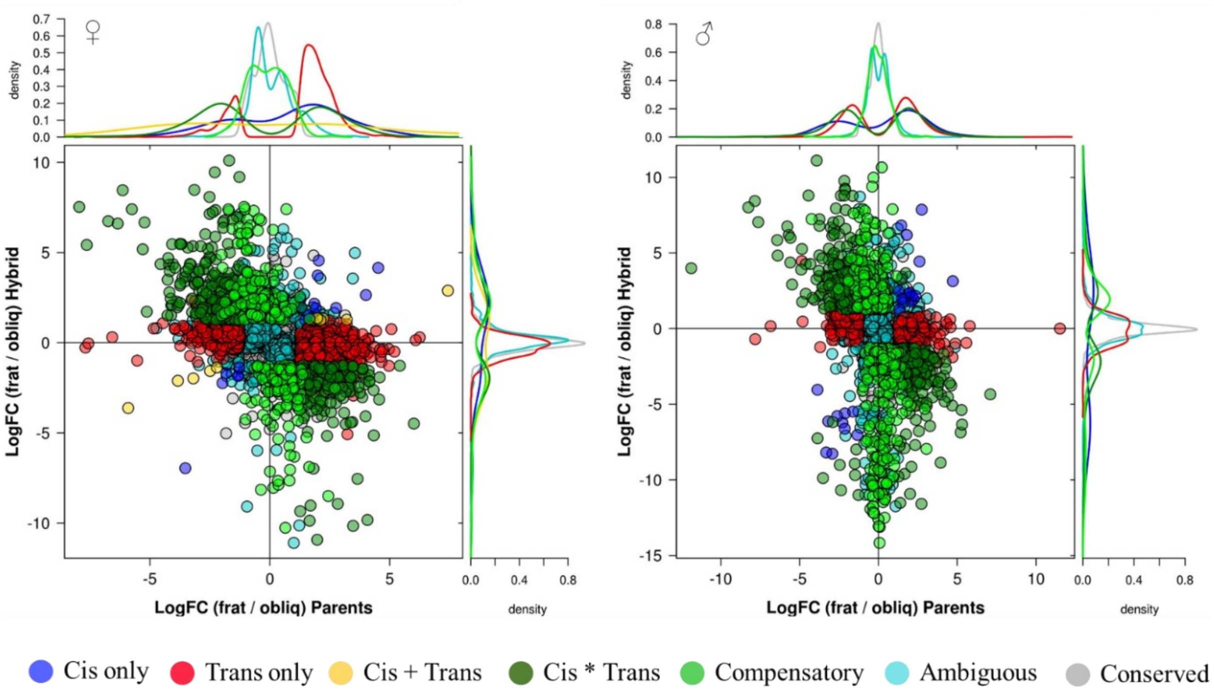


**Figure S3. Regulatory divergence due to combinations of *cis* and *trans* effects between *A. fraterculus* and *A. obliqua***. Scatterplots compare relative allelic expression levels between parental species [*$log_2$FC(frat/obliq)*] and between alleles in hybrids [*$log_2$FC(hyb_frat + hyb_obliq)*]. These results were used to sort genes into categories based on their mechanism of regulatory divergence.
